# DNA metabarcoding potentially reveals multi-assemblage eutrophication responses in an eastern North American river

**DOI:** 10.1101/186452

**Authors:** Joseph M. Craine, Matthew V. Cannon, Andrew J. Elmore, Steven M. Guinn, Noah Fierer

## Abstract

Freshwater aquatic ecosystems provide a wide range of ecosystem services, yet provision of these services is increasingly threatened by human activities. Directly quantifying freshwater biotic assemblages has long been a proxy for assessing changing environmental conditions, yet traditional aquatic biodiversity assessments are often time consuming, expensive, and limited to only certain habitats and certain taxa. Sequencing aquatic environmental DNA via metabarcoding has the potential to remedy these deficiencies. Such an approach could be used to quantify changes in the relative abundances of a broad suite of taxa along environmental gradients, providing data comparable to that obtained using more traditional bioassessment approaches. To determine the utility of metabarcoding for comprehensive aquatic biodiversity assessments, we sampled aquatic environmental DNA at 25 sites that spanned the full length of the Potomac River from its headwaters to the Potomac estuary. We measured dissolved nutrient concentrations and also sequenced amplified marker genes using primer pairs broadly targeting four taxonomic groups. The relative abundances of bacteria, phytoplankton, invertebrate, and vertebrate taxa were distinctly patterned along the river with significant differences in their abundances across headwaters, the main river, and the estuary. Within the main river, changes in the abundances of these broad taxonomic groups reflected either increasing river size or a higher degree of eutrophication. The larger and more eutrophic regions of the river were defined by high total dissolved phosphorus in the water, a unique suite of bacteria, phytoplankton such as species of the *diatom* Nitzschia, invertebrates like the freshwater snail *Physella acuta*, and high abundance of fish including the common carp (*Cyprinus carpio*). Taxonomic richness of phytoplankton and vertebrates increased downriver while it consistently decreased for bacteria. Given these results, multi-assemblage aquatic environmental DNA assessment of surface water quality is a viable tool for bioassessment. With minimal sampling effort, we were able to construct the equivalent of a freshwater water quality index, differentiate closely-related taxa, sample places where traditional monitoring would be difficult, quantify species that are difficult to detect with traditional techniques, and census taxa that are generally captured with more traditional bioassessment approaches. To realize the full potential of aquatic environmental DNA for bioassessment, research is still needed on primer development, a geographically broad set of reference sites need to be characterized, and reference libraries need to be further developed to improve taxonomic identification.

## Introduction

People rely on freshwater aquatic ecosystems for drinking water, recreation, fisheries, and agriculture. Yet, the quality of the water and the integrity of aquatic communities found in our lakes, rivers, and streams is increasingly threatened by human activities including agriculture, roads, industry, mining, human waste, urbanization, and deforestation [1, 2]. Poor water quality directly reduces quality of life and increases economic costs while reducing economic output [3]. Effective monitoring of water quality and the causes of water quality impairment is a critical step to maintaining our freshwater resources, preventing further degradation, and guiding restoration efforts. Although water quality can be measured directly, water quality can also be quantified through bioassessment—the utilization of species abundances to indicate environmental conditions [4, 5]. As opposed to direct measurements of environmental conditions, bioassessment of water quality provides the benefits of a more robust indicator of water characteristics and integrates over longer temporal and spatial scales than direct point measurements [6]. As different species differ in their responses to physical, biological, and chemical stresses and disturbances, bioassessment can indicate changes in a range of water quality metrics, including nutrients, pollutants, pH, clarity/turbidity, or temperature. Bioassessment uses the composition of biotic assemblages to infer stressors and disturbances with the assumption that individual taxa respond uniquely to these factors and the relative abundances of taxa can be used to infer the relative importance of individual stressors or disturbances [7].

Bioassessment typically involves the direct collection of organisms with their abundances quantified via visual inspection by trained taxonomists. For example, phytoplankton are typically identified and counted under a microscope to estimate the biovolume of different taxa in a given water sample [8]. Despite the widespread acceptance of these traditional bioassessment approaches [9], traditional visual assessment of the relative abundance of phytoplankton can be expensive, subject to observer bias that restricts comparisons over time and across observers, and constrained by low taxonomic resolution [10]. Macroinvertebrate assessment has similar constraints, but is further constrained by generally being limited to hard-bottom wadable streams [11]. Fish collection tends to be the most intensive sampling, is less effective for larger rivers than smaller rivers, and is less useful for those species that reside at depth or do not float when shocked [12, 13].

In contrast to traditional bioassessment, sequencing of aquatic environmental DNA (eDNA) via metabarcoding provides an alternate approach to assess the relative abundances of organisms in a given water body [14-16]. To accomplish this, DNA within a water sample is purified either directly from the water or from filtered particulates. Marker gene regions that are taxonomically informative (i.e. sufficiently variable in nucleotide composition to discriminate between different groups of organisms) are then amplified and sequenced, yielding information on the relative abundance of DNA from different organisms. Compared to traditional bioassessment approaches, sequencing aquatic eDNA is typically less expensive and often yields higher taxonomic resolution than visual assessment. Plus, sampling for eDNA analysis is often logistically easier than traditional assessment, can readily be performed in a wide range of different aquatic environments, and can be used to quantify abundances of organisms not traditionally censused. Although the eDNA approach is not bias-free and care must be taken when interpreting the results [17], the limitations of eDNA analysis are potentially offset by its advantages and the fact that the resulting data are not subject to observer bias, yielding datasets that are more reliable and consistent over time and space.

The utility of using eDNA-based metabarcoding to quantify aquatic organisms in rivers has already been demonstrated for some taxonomic groups [16, 18-21]. Despite the potential of metabarcoding for bioassessment, the technique has still not been tested extensively and we do not know whether multiple assemblages can simultaneously be assessed with current primer sets to generate biotic indices of environmental conditions. To examine the utility of aquatic eDNA metabarcoding for reconstruction of assemblages and bioassessment of environmental conditions, we sampled water from sites distributed along 475 km of the North Branch of the Potomac River from its headwaters to the Potomac estuary below Washington D.C. over a 3-day period in April 2017. The Potomac was chosen for assessment as it passes through a wide range of land uses from forest to agricultural to urban. Portions of the Potomac River are also considered to have experienced eutrophication due to agricultural and wastewater inputs, which in turn serve as a source of excessive nutrients for the Chesapeake Bay [22]. The Potomac is also the sole source of water for Washington D.C. and is an important recreational river for a large population. Given what is known about the Potomac, we employed metabarcoding to assess the relative abundances of bacteria, phytoplankton, invertebrates, and vertebrates using four primer pairs. At each site, we also analyzed water samples for total dissolved nitrogen and phosphorus as an independent estimate of nutrient availability in the water. We then assessed the multivariate correlations among the relative abundances of taxa and nutrient concentrations to test whether broad suites of taxa were responding similarly to changes in environmental conditions.

## Methods

### Site selection and sampling

Water samples were collected from 25 sites located on the North Branch or main stem of the Potomac River. These sites start at Fairfax Stone and end in the Potomac estuary downstream of Washington D.C., spanning a distance of 475 km. Among the 25 sites, 4 sites are considered headwaters (0-40 km), 9 sites were in the Upper Potomac (40-240 km), 9 sites were in the Lower Potomac (240-445 km), and 3 sites were in the Potomac estuary (>445 km downstream).

Sampling occurred between April 19-21, 2017. At each site, water was drawn into a sterile 60 mL syringe and then pushed through a Whatman Puradisc 25 mm 1µm nylon syringe filter. This process was repeated until no more water could be pushed through the syringe by the user. Across sites, an average of 252 mL was sampled, with a range of 120 to 500 mL of water sampled per site. The syringe filter was then placed into a 60 mL specimen cup with silica gel desiccant and stored at either room temperature or at 4°C until DNA was extracted.

### DNA sequencing

DNA was extracted from filters with a MoBio PowerSoil DNA kit (MoBio Laboratories, Carlsbad, CA) following the manufacturer’s protocol. For the phytoplankton analyses, we amplified a region of the 23S rRNA gene using PCR primers designed to amplify this gene region from a broad range of phytoplankton taxa, including Cyanophyta (cyanobacteria), Chlorophyta (green algae), and Bacillariophyta (diatoms) (Sherwood and Presting, 2007). Both primers also contained a 5’ adaptor sequence to allow for subsequent indexing and Illumina sequencing. Each DNA sample was amplified in triplicate reactions that were subsequently combined. These PCR reactions included 12.5 µL of Promega Mastermix, 0.5 µL of each primer, 1.0 µL of extracted DNA, and 10.5 µL of DNase/RNase-free H_2_O. The PCR reaction conditions consisted of an initial denaturation step of 3 min at 94°C, followed by 40 cycles at 94°C (30 seconds), 55°C (45 seconds), and 72°C (60 seconds), followed by a final elongation step of 10 minutes at 72°C. Similar procedures were used for bacteria, invertebrates, and vertebrates (Table 1).

**Table 1.**
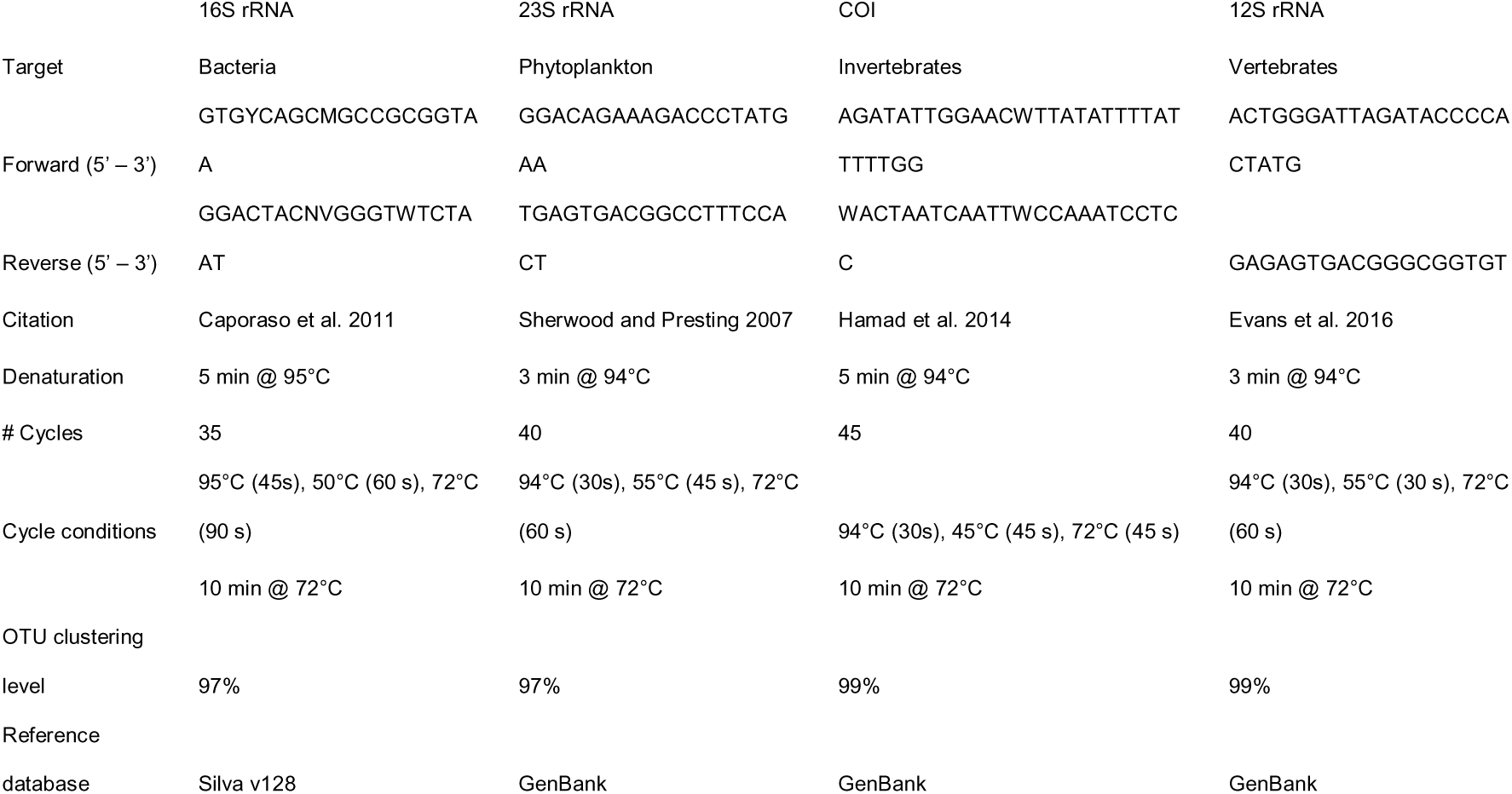
Details on unique laboratory methods and bioinformatic processing of data for different primer pairs.

After PCR, the amplicons were visualized on a 2% agarose gel to visually confirm that the PCRs yielded amplicons of the expected size. 20µl of the PCR amplicon was used for PCR clean-up using ExoI/SAP reaction. To index the amplicons with a unique identifier sequence, the first round of PCR was followed by an indexing 8-cycle PCR reaction to attach 10-bp error-correcting barcodes unique to each sample to the pooled amplicons from each site. These products were again visualized on a 2% agarose gel to check for band intensity and amplicon size. PCR products were purified and normalized using the Life Technologies SequalPrep Normalization kit and samples pooled together. Amplicons were sequenced on an Illumina MiSeq at the University of Colorado Next-Generation Sequencing Facility running the paired-end 2x250bp V2 sequencing chemistry.

### Bioinformatic processing

After de-multiplexing the reads, the paired-end reads were merged using fastq_merge pairs [23]. Since merged reads often extended beyond the amplicon region of the sequencing construct, we used fastx_clipper to trim primer and adapter regions from both ends (https://github.com/agordon/fastx_toolkit). Sequences lacking a primer region on both ends of the merged reads were discarded. Sequences were quality trimmed to have a maximum expected number of errors per read of less than 0.1 and only sequences with more than 3 identical replicates were included in downstream analyses. BLASTN 2.2.30+ was run locally, with a representative sequence for each operational taxonomic unit (OTU) as the query and the current National Center for Biotechnology Information (NCBI) nt nucleotide and taxonomy database as the reference. The tabular BLAST hit tables for each OTU representative were then parsed so only hits with > 97% query coverage and identity were kept. Similar procedures were used for the other primer pairs (Table 1).

The 23S rRNA sequences were clustered into OTUs at the ≥97% sequence similarity level and sequence abundance counts for each OTU were determined using the usearch7 approach. The NCBI genus names associated with each hit were used to populate the OTU taxonomy assignment lists. Sequences that did not match over 90% of the query length and did not have at least 85% identity were considered unclassified, otherwise the top BLASTn hit was used for taxonomy assignment. Similar procedures were used for the other primer pairs (Table 1).

For the 23S rRNA gene analyses (phytoplankton), we removed all OTUs that were identified as higher plants or uncultured organisms. The average number of remaining reads per sample was 12454. For the 12S rRNA gene analyses (vertebrates), we removed all taxa except those assigned to Chordata. The average number of remaining reads for 12S rRNA was 1096 per sample. For the COI gene analyses (macroinvertebrates), we removed all OTUs except those assigned to Annelida, Arthropoda, Cnidaria, and Mollusca. After removing all other taxa, the average number of remaining COI reads per samples was 125. The reason there were so few reads was that >95% of the reads were from Oomycota (3% were from Rotifera). The COI gene from oomycetes, i.e. water molds, is amplified with the primers we used and oomycetes happen to be highly abundant in the Potomac. As our target taxa were macroinvertebrates, we had made the decision before analyzing the data to exclude all taxa that were not macroinvertebrates and did not include oomycetes in our analyses here.

Taxonomic identifications were constrained by the availability of sequences in our reference databases. As such, some OTUs were likely assigned to species that were closely related, but not identical, to those present in the Potomac. For example, one OTU was assigned to *Cottus szanaga*, which is only found in Asia. No other *Cottus* species were identified. More than likely, the DNA was derived from a different *Cottus* species present in the Potomac that has not been sequenced yet such as *Cottus caeruleomentum* or *Cottus girardi*. In all cases, we refer to OTUs based on the species to which they were matched despite these limitations.

### Nutrient analyses

At each site, 250 mL of filtered water was retained in a sterile scintillation vial and kept cold and in the dark. In the laboratory, the water was re-filtered with a 0.45 µm filter and frozen at -20°C for analysis within 28 d. Thawed water was analyzed for total dissolved phosphorus and nitrogen at the University of Maryland Center for Environmental Science, Appalachian Laboratory’s Water Chemistry Analytical Lab using offline persulfate digestion followed by colorimetric analysis for orthophosphate and nitrate+nitrite on a Lachat QuikChem 8000 Flow Injection Analyzer.

### Statistical analyses

With 3-5 lab replicates for each sample, all data for the lab replicates were summed for each sample. With multiple OTUs often assigned to the same taxon, all data on the number of reads was summed for vertebrate, macroinvertebrate, and phytoplankton data based on taxonomic identity. Bacterial 16S rRNA data were retained at the OTU level.

To estimate taxonomic richness, the number of reads for each sample was rarefied to a set number of randomly selected reads per sample to control for differences in sequencing depth. 16S rRNA data were rarefied to 22698 reads per sample, 23S rRNA data were rarefied to 6564 reads, 12S rRNA data to 762 reads. Rarefied richness was not calculated for COI data due to there being too few reads per samples with these primers.

To assess the general patterns of the abundances of taxa with equal weighting among the major taxonomic groups, we ran principal components analyses (PCA) with the top 30 taxa for each primer pair. We also ran a single PCA that included the top 30 taxa for each primer pair together as well as rarefied richness for taxa identified with 16S rRNA, 23S rRNA, and 12S rRNA primer pairs.

Sørensen’s index of similarity was calculated for all pairs of samples for each of four groups of taxa using the *betadiver* command of *vegan* package in R. Similarity indices between sampling points were compared with hydrologic distances along the river with a Mantel test from the *vegan* package [24]. *P* values for the index of similarity were calculated as 1-*p* where *p* is the likelihood of a randomization permutation resulting in a matrix becoming more dissimilar with hydrologic distance. For each taxon, we also calculated the average distance each taxon was found along the river by calculating the average distance of all samples weighted by the relative read abundance of that taxon.

All analyses were conducted in R version 3.3.2 except for the PCAs, which were conducted in JMP 13.0.0 (SAS Institute Inc., Cary, NC, USA).

## Results

### Taxa-level patterns

For bacteria sampled across the Potomac, 52% of the reads were assigned to Proteobacteria, 22% to Bacteroidetes, 14% to Actinobacteria, and 6% to Verrucomicrobia. Proteobacteria averaged 60% of the reads through 300 km and then declined in the Lower Potomac to as low as 30%. In contrast, Bacteriodetes, Actinobacteria, and Verrucomicrobia all increased in relative abundance with increasing distance downstream (*P* < 0.001 for all), peaking just before the estuary. Individual bacterial OTUs were uniquely distributed in ways that mirrored phylum-level patterns, but not always. For example, the most abundant bacterial OTU (OTU2, Proteobacteria, Sphingomonadales) was found on average 275 km downstream, while another Proteobacteria (OTU 25, Hyphomonadaceae) was found on average 114 km downstream. Average rarefied richness declined with distance downstream (*P* < 0.001), declining at a rate of 2.66 ± 0.49 OTUs km^-1^.

Across the phytoplankton dataset, 78% of the reads were assigned to Bacillariophyta, 3.4% to Eustigmatophyta, 2.0% to Cyanobacteria, and 1.8% to Chlorophyta. Diatom (Bacillariophyta) read abundance was ∼20% for the first 10 km and then increased in abundance with distance downriver, dominating the rest of the river and estuary, representing 85% of the reads, on average, after 50 km (Figure 2). Examining the abundance-weighted distances of all diatom reads, diatoms were most abundant 252 km downstream. In contrast, eustigmatophytes dominated reads from the headwaters, with a given eustigmatophyte located on average at just 59 km downstream (Figure 2).

**Figure 1.**
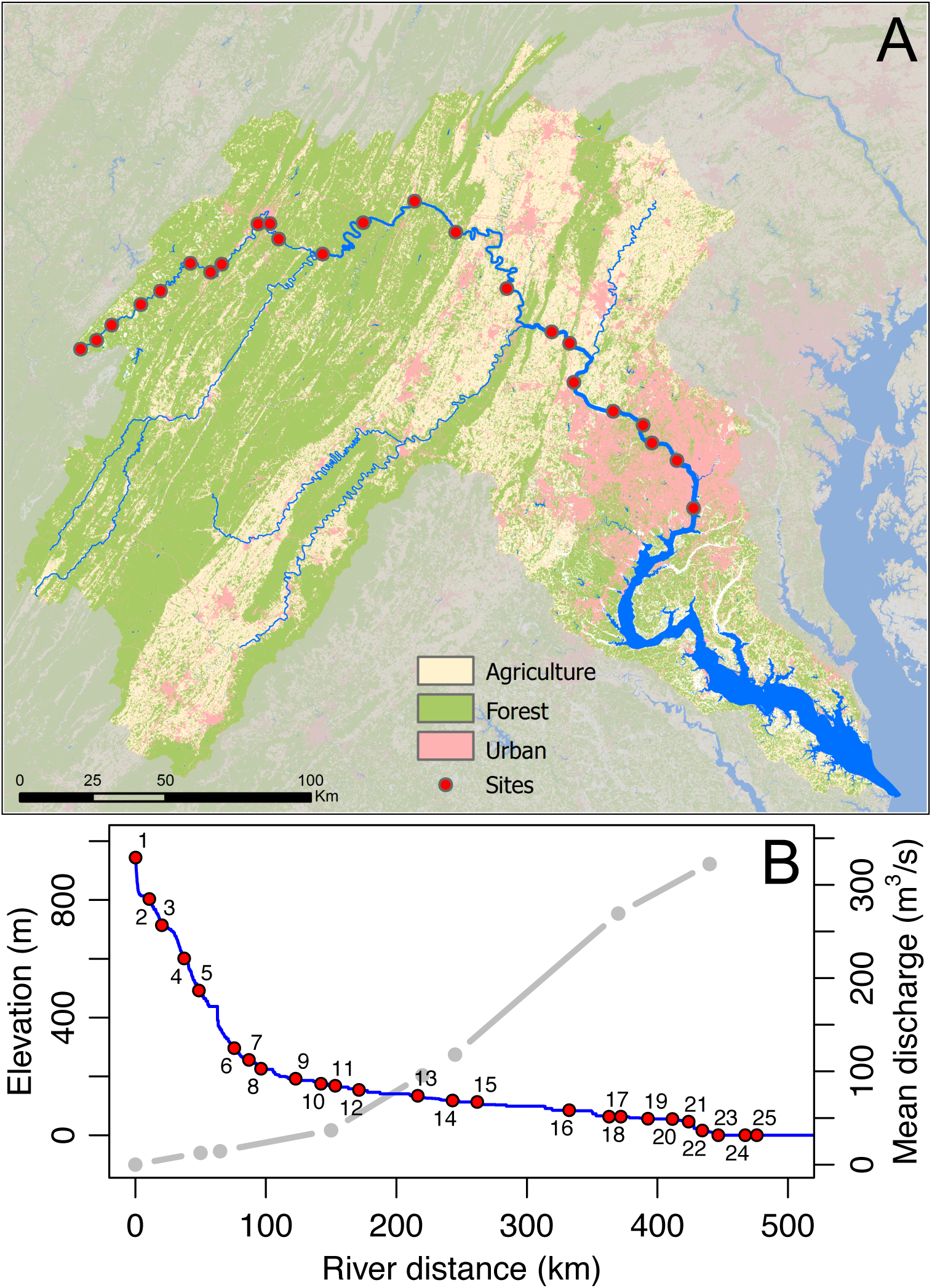
Map of the locations of the 25 sites sampled for this study (A), the elevation of each site (B), and mean annual discharge for selected points from USGS streamflow data (B).

**Figure 2.**
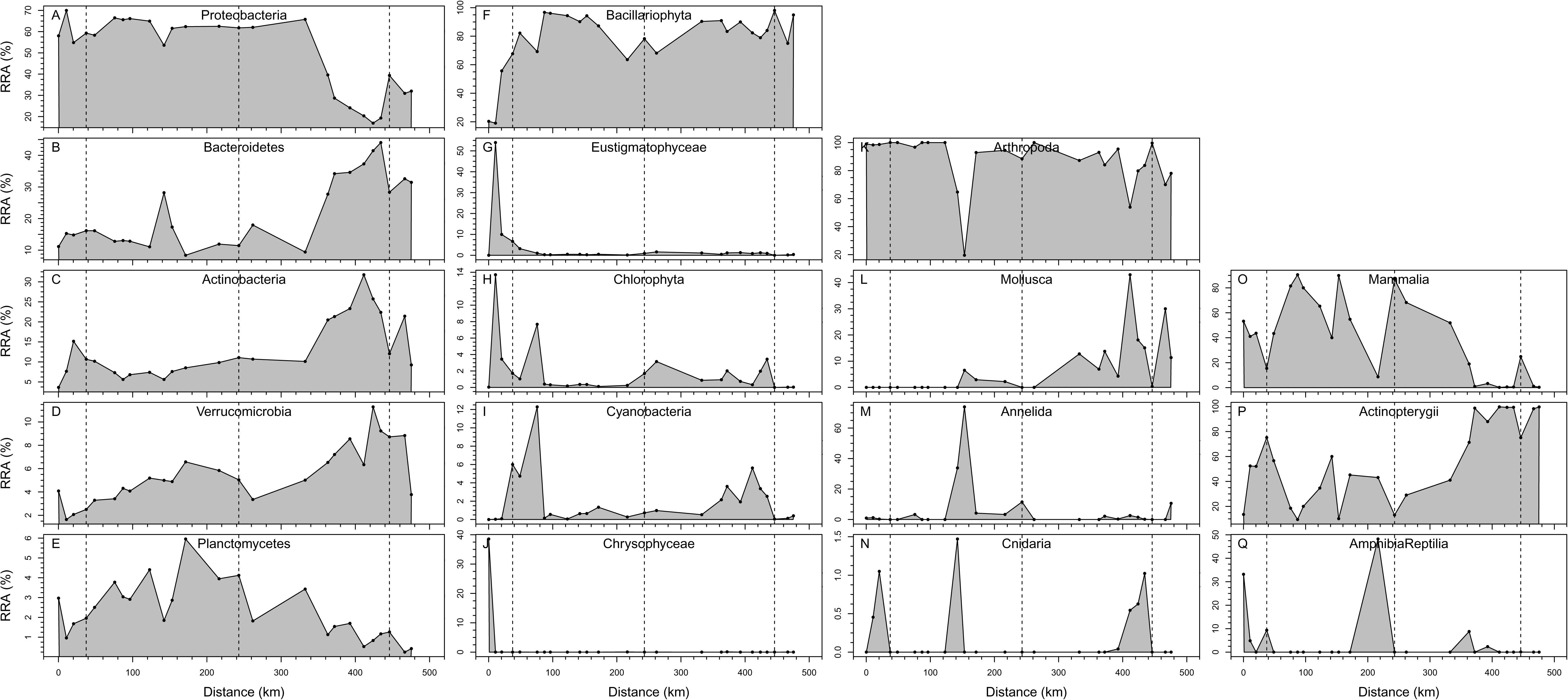
Percent relative read abundance (%RRA) of sequences assigned to higher order taxonomic groups as a function of distance downriver. Included are abundances for bacteria (A-E), phytoplankton (F-J), macroinvertebrates (K-N), and vertebrates (O-Q) primers.

Individual phytoplankton taxa revealed distinct patterning along the Potomac. For example, the most abundant phytoplankton taxon on average was the diatom *Navicula salinicola,* and was most abundant in the upper estuary (Figure 3). On average, it was located 367 km downriver. In contrast, the diatom Nitzschia sp. [BOLD:AAX5147] was the third most abundant phytoplankton and more abundant in Upper Potomac, found on average 148 km downriver (Figure 3). Observed phytoplankton species of general interest included *Didymosphenia geminate,* a.k.a. “rock snot”, which can form large mats in nutrient-poor waters and can bloom to nuisance levels. It was found on average 103 km downriver where it accounted for >10% of the phytoplankton sequences, with the highest relative abundance in the middle of the sampled reach. Cyanobacteria species of the genus *Synechococcus* were more abundant in the lower Potomac, but were relatively rare (< 5% of the reads). Rarefied richness of phytoplankton taxa increased with increasing distance and dropped dramatically in the estuary (Figure 4).

**Figure 3.**
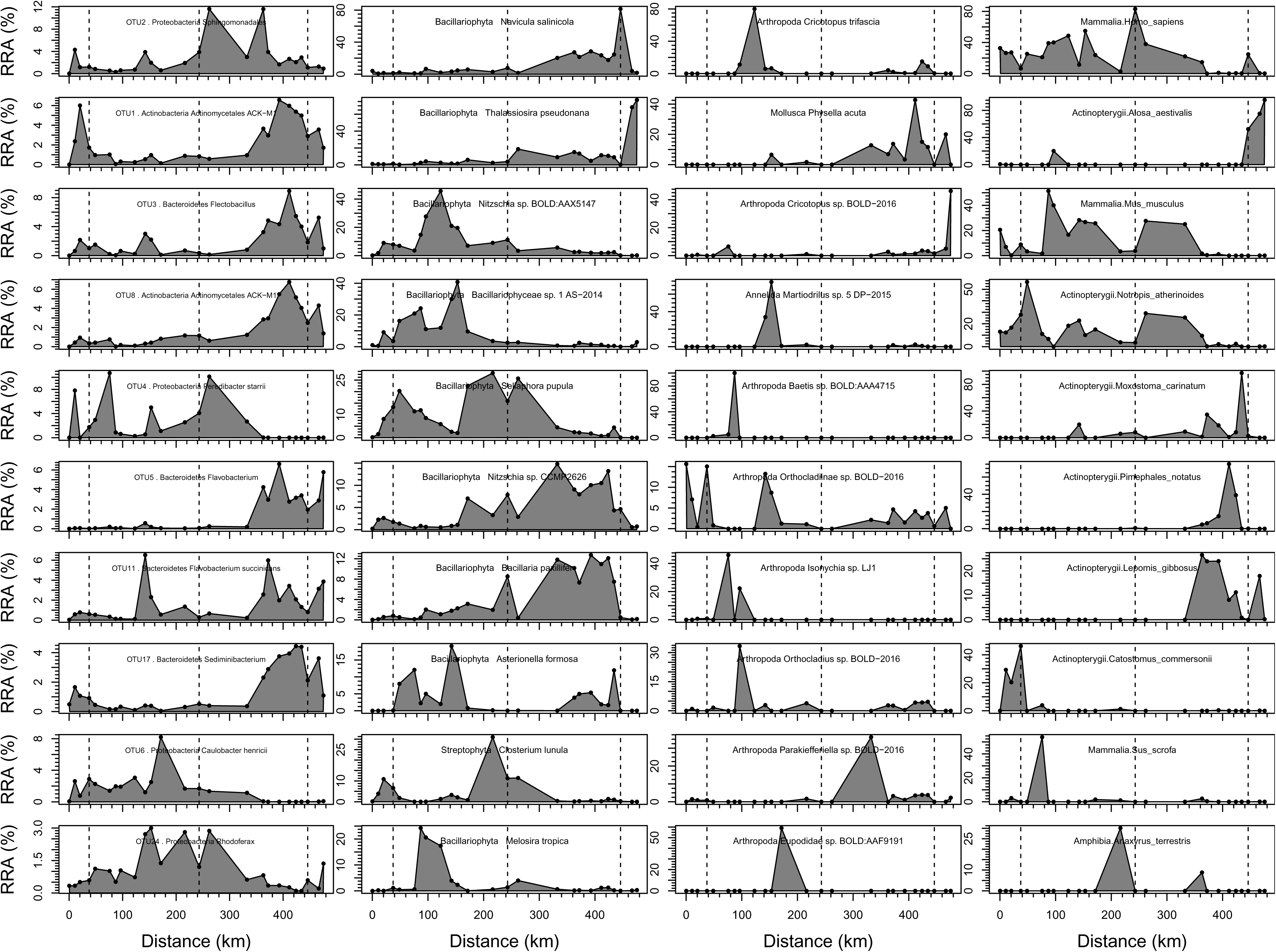
Percent relative read abundance (%RRA) of the ten most abundant taxa for bacteria (first column), phytoplankton (second column), macroinvertebrates (third column), and vertebrates (fourth column) primers plotted versus distance downriver.

For the macroinvertebrates identified via COI sequencing, 87% of the reads were assigned to Arthropoda, 7% were to Mollusca, and 6% to Annelida. Of the reads assigned to Arthropoda, 61% were Diptera, 20% Ephemeroptera, and 5% Plecoptera. Like the phytoplankton, invertebrate taxa were differentially abundant along the Potomac. For example, the most abundant invertebrate taxa, the Diptera *Cricotopus trifascia*, was found towards the middle region of the Potomac, while the second most abundant invertebrate taxa, the mollusk *Physella acuta* was found closest to the mouth of the Potomac (Figure 3).

Of the 80 vertebrate taxa identified, 39% of the reads were Mammalia, including sequences identified to humans, mice, pigs, cattle, white-tailed deer, beaver, raccoon, bank voles, and muskrat. 56% of the reads were Actinopterygii including sculpin (*Cottus*), darters (*Etheostoma*), shad (*Alosa*), sunfish (*Lepomis*), suckers (*Catostomus*), trout (*Salmo* and *Oncorhynchus*), and carp (*Cyprinus carpio*). 5% of the reads were assigned to Amphibia and Reptilia, including the American toad (*Anaxyrus americanus*), the American bullfrog (*Rana catesbeiana*), snapping turtle (*Chelydra serpentina*), and the northern dusky salamander (*Desmognathus fuscus*). Human DNA was the most abundant vertebrate DNA recovered from the water samples, averaging 21.7% of the 12S rRNA gene reads with human DNA generally most abundant in the upper Potomac (Figure 3). The second most abundant vertebrate taxon identified was *Alosa aestivalis*, but likely represented multiple *Alosa* species, if not related species. Approximately 10% of the reads were assigned to *Alosa aestivalis* and this taxon was dominant in the estuarine samples (*P* < 0.001; Figure 3). When all samples were rarefied to 762 reads, vertebrate diversity averaged 11.5 species and peaked at 25 species ∼390 km downriver (Figure 4).

**Figure 4.**
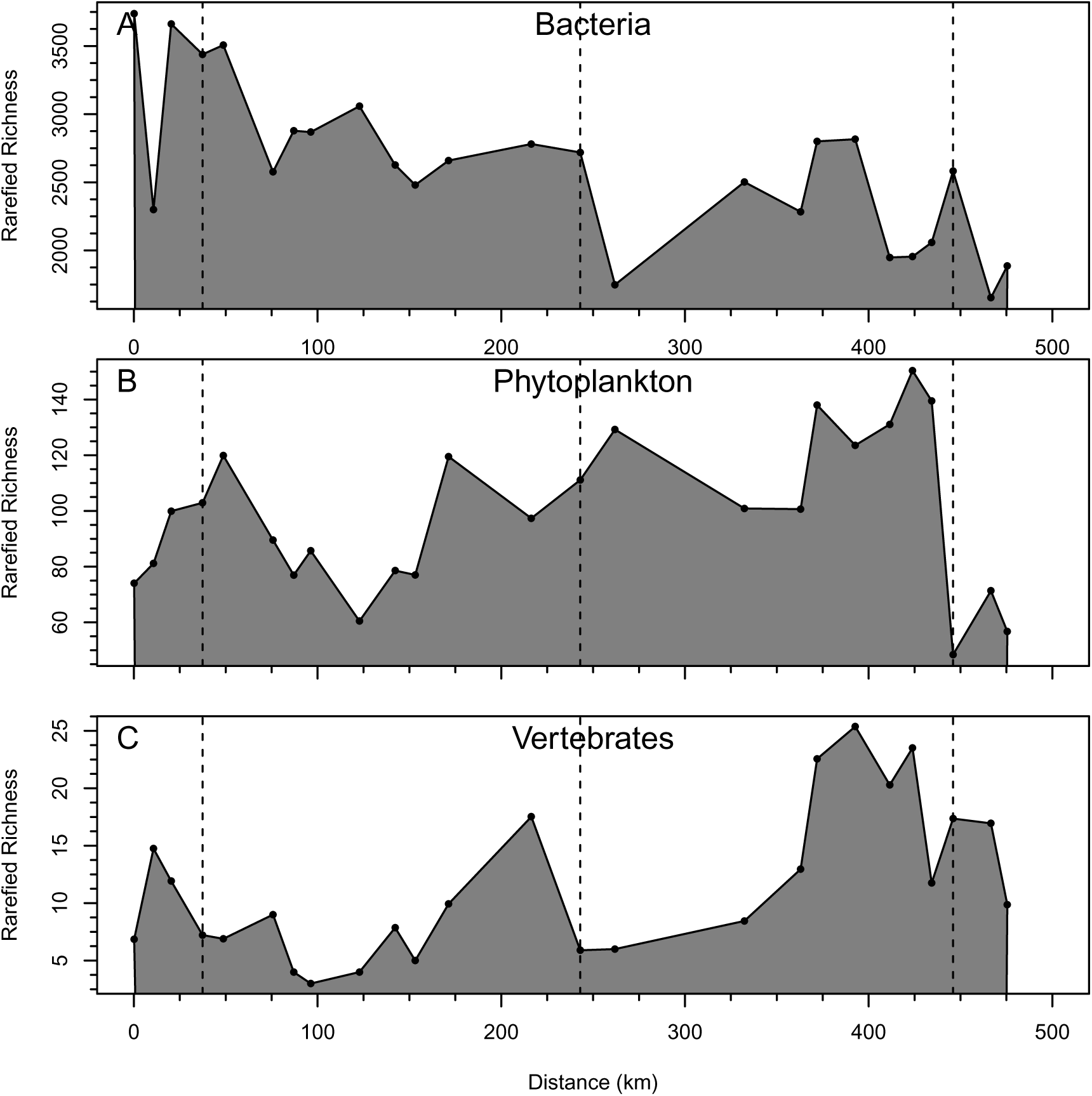
Rarefied richness of taxa as a function of distance downriver for bacteria (A), phytoplankton (b), and vertebrates (c).

Assessing the Sørensen’s index of community similarity as a function of distance for each taxonomic group, adjacent samples were projected to have similarity values of 55.7% for bacteria, 65.3% for phytoplankton, 21% for invertebrates, 50.1% for vertebrates. Similarity declined with distance for all four groups (*P* < 0.001; Figure 5). The rate of decline in similarity was greatest for vertebrates (7.1% 100km^-1^); intermediate for bacteria and phytoplankton (4.7% and 4.6% 100km^-1^), and lowest for invertebrates (2.6% 100km^-1^), which had the lowest degree of similarity for geographically adjacent assemblages.

**Figure 5.**
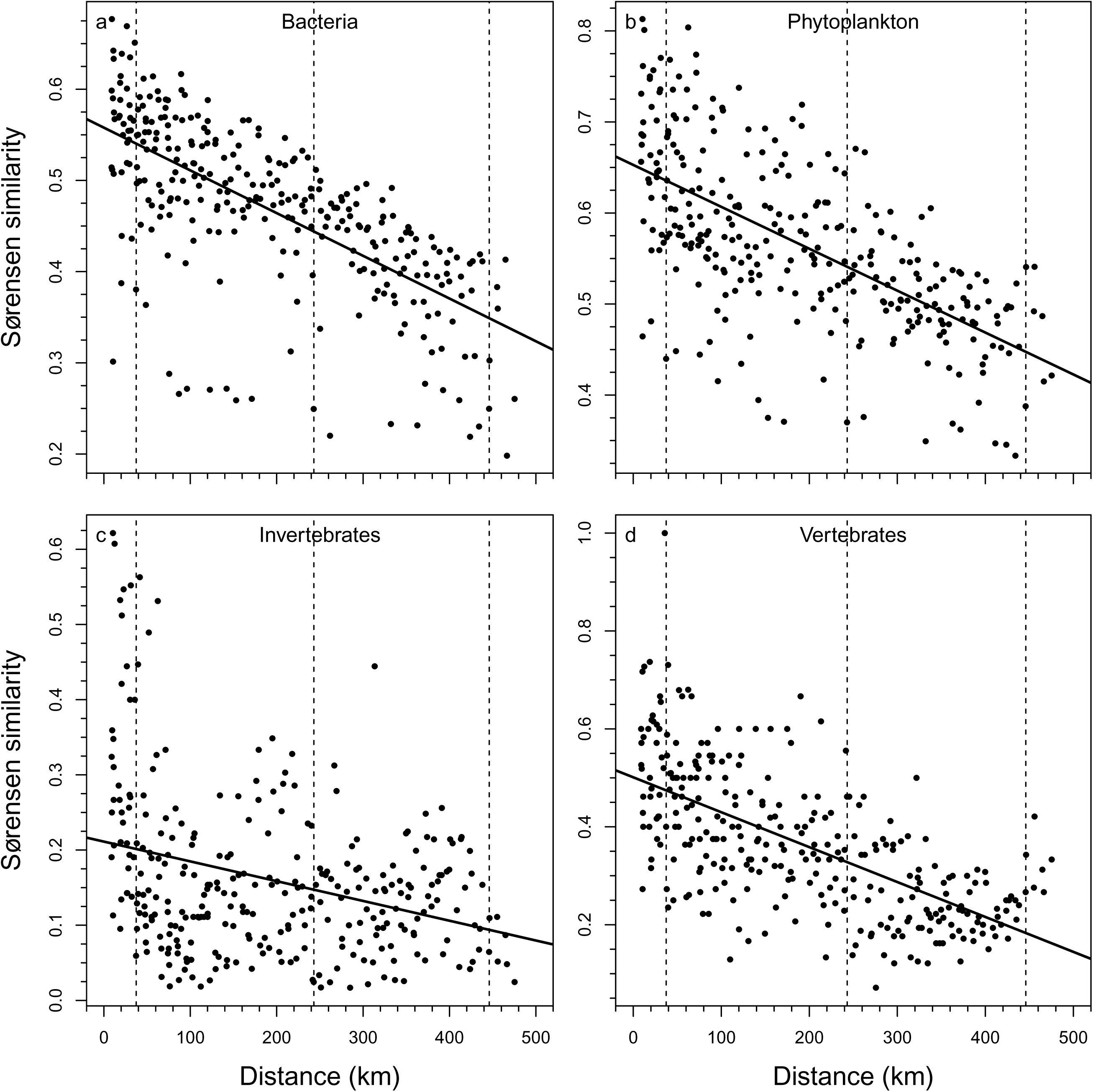
Sørensen’s index of similarity for bacteria (A), phytoplankton (B), invertebrates (C), and vertebrates (D) as a function of distance between 2 of the 25 water samples collected on the Potomac River. Lower values indicate communities that are more distinct in composition. Best fit linear regressions shown with Mantel test used to assess significance of correlation (ρ = -0.67, -0.64, -0.32, -0.64, respectively; *P* < 0.001 for all).

### Nutrients

As with the biotic communities, nutrient concentrations also showed distinct patterns with distance downriver (Figure 6). Total dissolved phosphorus concentrations were generally the lowest in the headwaters and gradually increased until peaking in the Lower Potomac. In contrast, total dissolved nitrate concentrations were high in the headwaters, were lowest in the Upper Potomac, and were high in Lower Potomac and estuary samples (Figure 6).

**Figure 6.**
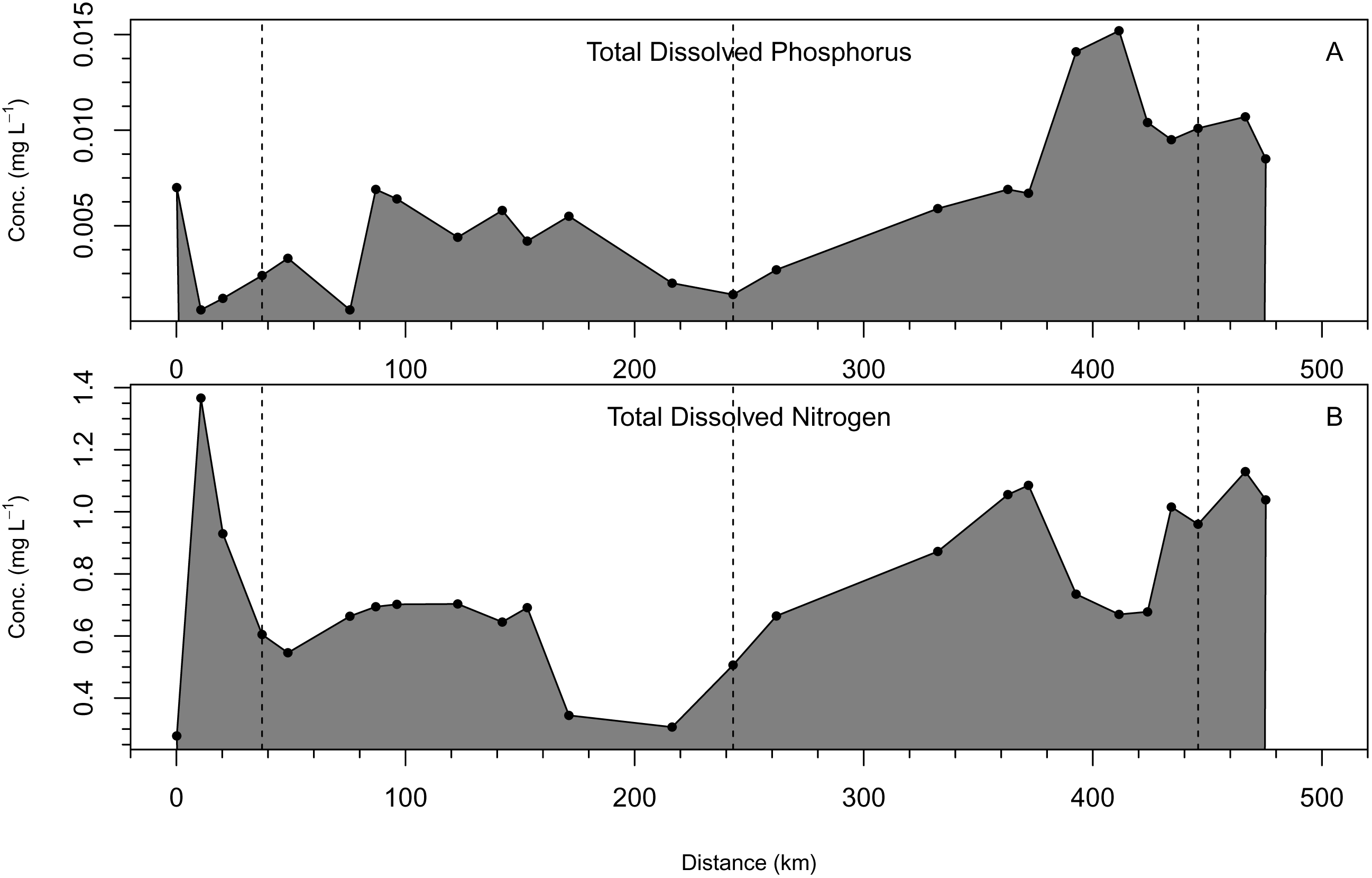
Availability of phosphorus (A) and nitrogen (B) as a function of distance on the Potomac River.

### Multivariate patterns in overall assemblage composition

We examined multivariate patterns of relative abundance with principal component analysis to assess the broad patterns of changes in relative abundance among taxa without *a priori* assumptions of what factors might be structuring the observed patterns. Examining the results of the principal component analysis for each primer pair, the first multivariate axis in each PCA appeared to capture either increasing phosphorus availability, i.e. eutrophication, or increasing river size except for COI (Figure 7). A PCA analysis with all surveyed taxa combined revealed a stronger pattern of shifts in taxa abundances with either downstream eutrophication or river size. In the multi-assemblage PCA, Axis 1 explained 20.1% of the data, which is 23.4 times more than expected by chance. Axis 1 increased with river distance peaking just before Great Falls where the river becomes tidal (Figure 7). Axis 1 correlated with distance (r = 0.85, *P* < 0.001) and total dissolved phosphorus (r = 0.78, *P* < 0.001), but not total dissolved nitrogen (*P* = 0.09).

**Figure 7.**
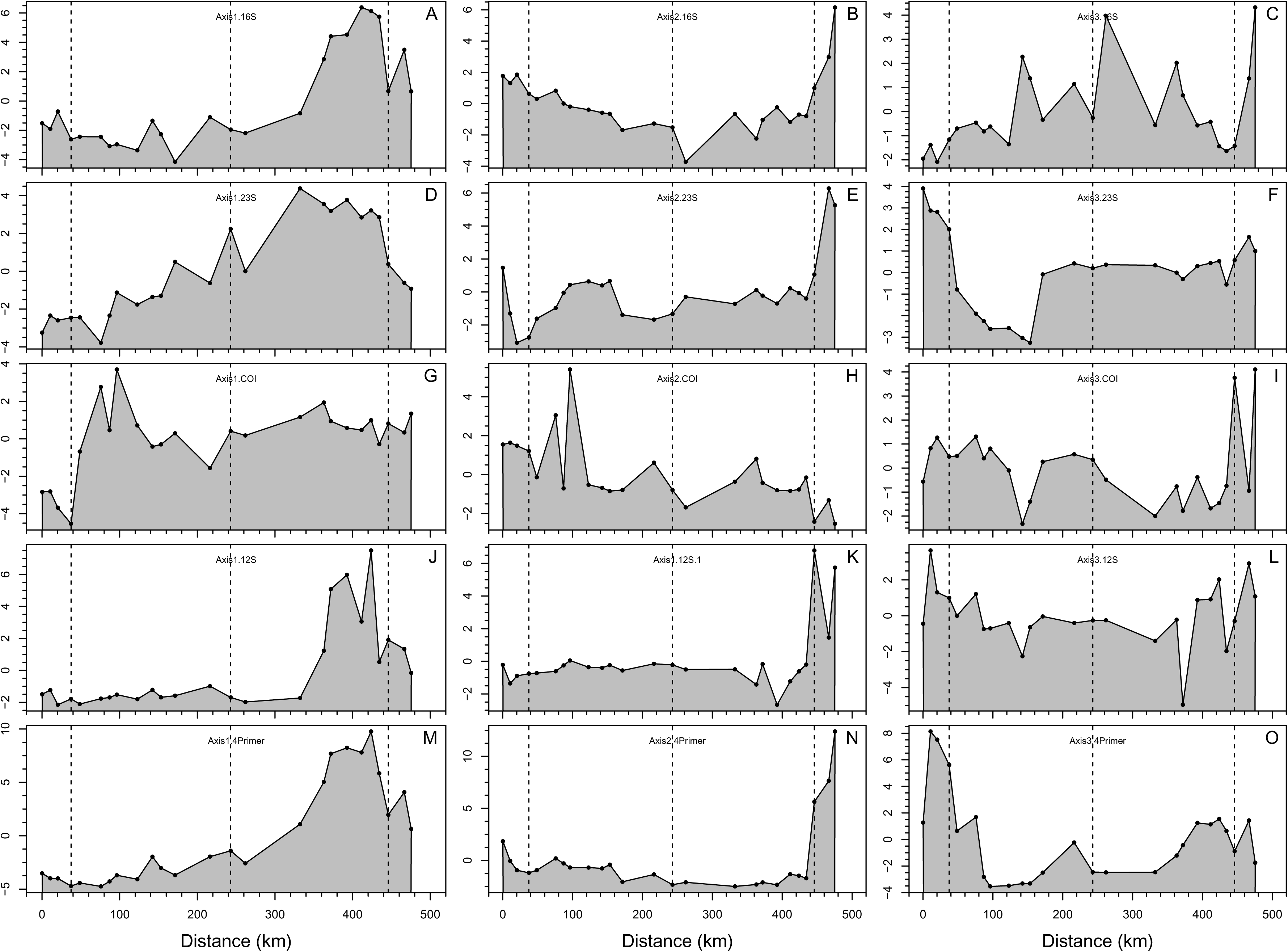
Scores of sites on first three axes of principal components analyses (PCA) for bacteria (A-C), phytoplankton (D-F), macroinvertebrates (G-I), vertebrates (J-L), and all 4 primer pairs together (M-O). The specific y-axis scores simply represent the position of each assemblage along each individual PCA axis with assemblages that are more similar in composition having more similar axis scores.

We next examined the coefficients of the variables in the PCA to assess which taxa were the most important contributors to the general patterns described above. Of the top 10 taxa with the highest coefficients for the multi-assemblage Axis 1, 8 were bacteria (Table S1). These included the Actinomycete of ACK-M1 group and the Bacteriodetes *Sedimentibacteria*. The other taxa were the common carp (*Cyprinus carpio*) and a diatom indicative of eutrophic conditions (*Bacillaria paxillifer*). Other select taxa with high Axis 1 coefficients include redspotted sunfish (*Lepomis miniatus*), spotfin shiner (*Cyprinella spiloptera)*, which is often found in poor quality waters, a mayfly that tends to inhabit large rivers, *Anthopotamus verticis*, the snail *Physella acuta*, which often is found in degraded waters, and a diatom *Nitzschia* indicative of eutrophic waters.

Axis 1 also distinguished among taxa that were found in higher abundance in the Upper Potomac, but not exclusively in the headwaters (Table S1). Of the 10 taxa with the lowest coefficients on Axis 1, five were Proteobacteria including a Hyphomonadaceae OTU and a Rhodobacter OTU. Other taxa included the diatoms *Didymosphenia geminata* and *Sellaphora pupula* and fish species similar to the emerald shiner (*Notropis atherinoides)* (Table S1). Human DNA was also more abundant in Upper Potomac and had a low coefficient on Axis 1 (Table S1). Although many of the strongest taxa driving the separation of the communities along Axis 1 were associated with bacteria, the scores of sites on a separate PCA with the phytoplankton, macroinvertebrate, and vertebrate taxa, but without bacteria, were strongly correlated with those generated with all four sets of taxa (*r* = 0.96, *P* < 0.001).

The second axis of the multi-assemblage PCA primarily separated out taxa that were more abundant in the estuary (Figure 7). Bacteria that were more abundant in the estuary included multiple cyanobacterial OTUs. Phytoplankton that were more abundant in the estuary were diatoms such as *Cyclotella* sp. [WC03_2] and taxa similar to *Thalassiosira pseudonana*, a marine diatom (Table S1). Invertebrates that were relatively abundant included a *Cricotopus* species (a dipteran) and the Asian clam, *Corbicula fluminea*, which accounted for >10% of the reads in the most downstream site (Table 1). Fish such as white perch (*Morone americana*), american eel (*Anguilla rostrata*), shad (*Alosa aestivalis*), and gizzard shad (*Dorosoma cepedianum*) were dominant in the estuarine samples (Table 1, Supplemental).

Axis 3 primarily distinguished headwater sites, which harbored a unique set of taxa (Table 1). Headwater bacteria included the Proteobacteria *Mycoplana* and *Rhodobacter*. Headwater phytoplankton species were generally taxa other than diatoms such as the eustigmatophyte, *Nannochloropsis salina,* the chlorophyte *Choricystis parasitica,* and the green alga, *Actinotaenium cruciferum* (Table S1). Arthropods included the mayfly, *Ephemerella dorothea* and the Diptera *Parametriocnemus* sp. BOLD-2016 (Table S1). Fish characteristic of the headwater samples included the white sucker (*Catostomus commersonii*), and sculpin (identified as *Cottus szanaga*).

## Discussion

By analyzing eDNA collected from the Potomac River, we were able to demonstrate the potential utility of eDNA for assessing aquatic assemblages and constructing multi-taxa indices that reflect environmental conditions. Here, using four primer pairs, we were able to quantify the DNA of resident bacteria, phytoplankton, invertebrates, and vertebrates across the length of a major river. Although better reference databases are needed to improve taxonomic identifications, the taxa observed corresponded well to taxa observed in previous surveys of rivers in the region [25-28]. Abundant bacterial taxa observed here, e.g. *Sphingomonodales*, *Flectobacillus*, and *Flavobacterium succinicans*) are typical of rivers [27, 29, 30]. Previous work has shown that riverine and estuarine phytoplankton communities share many of the same taxa observed here, e.g. *Navicula*, *Thalassiosira*, and *Nitzschia* [31-33]. Likewise, the more abundant invertebrate and vertebrate species we detected (Figure 1) are taxa we would expect in these types of river systems [25, 34-36].

In addition to identifying the composition of the biotic assemblages, we were able to assess correlations in taxon abundances across sites. We found that the biotic assemblages were spatially patterned and reflected changing environmental conditions across the length of the Potomac River. We were also able to clearly delineate changes in assemblages across the headwaters, the main river, and the estuary with the strongest shift in biotic assemblages observed between the headwaters and the estuary. Most likely, this gradient represents some combination of taxa responding to eutrophication and river size, which could not cleanly be separated here given the general increases in phosphorus concentrations with distance downstream. Along this gradient, the river increased in size from a small spring with a discharge of <1 m^3^ s^-1^ at the Fairfax Stone, to a mean discharge of 322 m^3^ s^-1^ at 440 km (Little Falls). We found stronger correlations between assemblage composition and distance along the river than with total dissolved phosphorus. Although phosphorus availability was measured with spring baseflows, which are considered good indicators of annual discharge-weighted mean concentrations [37], nutrient concentrations were single point measurements and might not necessarily represent integrated annual availability, to which the organisms are more likely responding. Alternatively, the observed patterns may be driven by differences in eutrophication levels with distance along the river, an interpretation supported by the observation that many of the taxa driving the patterns in assemblage composition evident in Figure 7 are taxa frequently associated with eutrophication status. Many *Nitzschia* and *Bacillaria* taxa are considered indicators of eutrophication [38, 39]. Fish such as *Cyprinus carpio* and *Cyprinella spiloptera,* as well as the snail *Physella acuta* are also associated with waters degraded to different degrees [40-42]. Many of the taxa that scored low on Axis 1 are associated with low nutrient conditions, such as the diatom *Didymosphenia geminate* [43, 44]. Yet, in contrast, one mayfly taxa, *Anthopotamus verticis,* was associated with high Axis 1 scores. Most mayfly species are considered indicators of high, not low, water quality, but *Anthopotamus* species often occupy waters with intermediate rather than low P availability [45].

Despite the relative success at this attempt to generate multi-assemblage indices of environmental conditions, more research and development is needed in a number of areas before the use of environmental DNA for bioassessment can be widely adopted. The uncertainty of the interpretation of assemblage compositions highlights the need for a comprehensive process to calibrate indices generated with eDNA. Multiple reference sites will need to be established that allow for separation of covariates such as river size and eutrophication. For example, multiple large rivers of different nutrient status will have to be sampled to separate out river size and nutrient conditions as drivers of taxon abundances. River systems that span gradients in other environmental variables, such as pH and temperature, will have to be surveyed to further separate out other covariates. More research on individual taxa will also be needed to link taxon abundances to environmental conditions. Although indices can be generated independent of taxonomy [46], building ecological understanding of individual taxa will assist in interpreting multitaxa indices of environmental conditions.

Our results also highlight the need for the development of more effective primer pairs. To assess macroinvertebrates (namely arthropods), we used a primer pair that is considered to be relatively specific for arthropods [47]. However, 95% of the reads with these COI primers were assigned to Oomycota taxa. Oomycetes are commonly known as water molds, include plant and animal pathogens as well as taxa that are important decomposers of organic matter in water [48, 49]. A full assessment of the patterns of Oomycota in the Potomac was outside the scope of this research, but is warranted given their importance in ecosystem function and their potential roles as pathogens. In the future, assessing invertebrates in water using eDNA will either require greater sequencing depth, blocking primers, or the development of different primers that are more selective for arthropods. As an additional note, we also observed relatively few vertebrate reads per sample, but this result was likely due to interactions during sequencing that disfavor longer 12S rRNA amplicons when multiplexed with amplicons generated with other primers. Sequencing only 12S rRNA amplicons on a given sequencing run should relieve this limitation.

The benefits of using eDNA to reconstruct assemblages appear to extend beyond the ability to reconstruct of environmental conditions. For example, as in a previous study of the Cuyahoga River [21], we were able to detect different amounts of mammalian DNA (including human, pig, and cattle DNA) with these results representing a potential opportunity to identify sources of fecal contamination in water. Current approaches often rely on the bacteria that indicate vertebrate sources of fecal coliform bacteria [50]. Yet, it might be feasible to simply sequence host DNA directly from collected water samples. Sequencing eDNA also has the potential to identify non-native and migratory species [51]. Here, we detected migratory fish such as shad and eel in the estuary, although some species known to be in the Potomac, such as northern snakehead (*Channa argus*), were not detected here. Greater sequencing depth and replication would likely provider greater sensitivity than our preliminary assessment, but probabilities of detection have yet to be resolved for individual taxa of interest.

Using eDNA to reconstruct assemblages also opens a new line of research for development of ecological theory. Although the river continuum concept was a crucial development for beginning to understand changes in ecosystem function along the length of a river [52, 53], theoretical development of the regulation of biodiversity within rivers has lagged [54, 55]. Data here raise interesting questions. For example, why did bacterial diversity decline downriver while it increased for phytoplankton and vertebrates? Are bacterial assemblages being driven by environmental chemistry as observed in other rivers [56]? Questions about accumulation of species, changes in habitats, nestedness of assemblages, and assemblage turnover are all relevant to understanding the ecologies of river systems that could be addressed by leveraging eDNA approaches to obtain large amounts of relatively low-cost standardized data for rivers.

## Acknowledgments

Amy Beckley carried out the lab work for this research, Mike Robeson was responsible for bioinformatics. Kristy Deiner provided helpful comments on previous drafts of this manuscript.

---

**Table S1.**
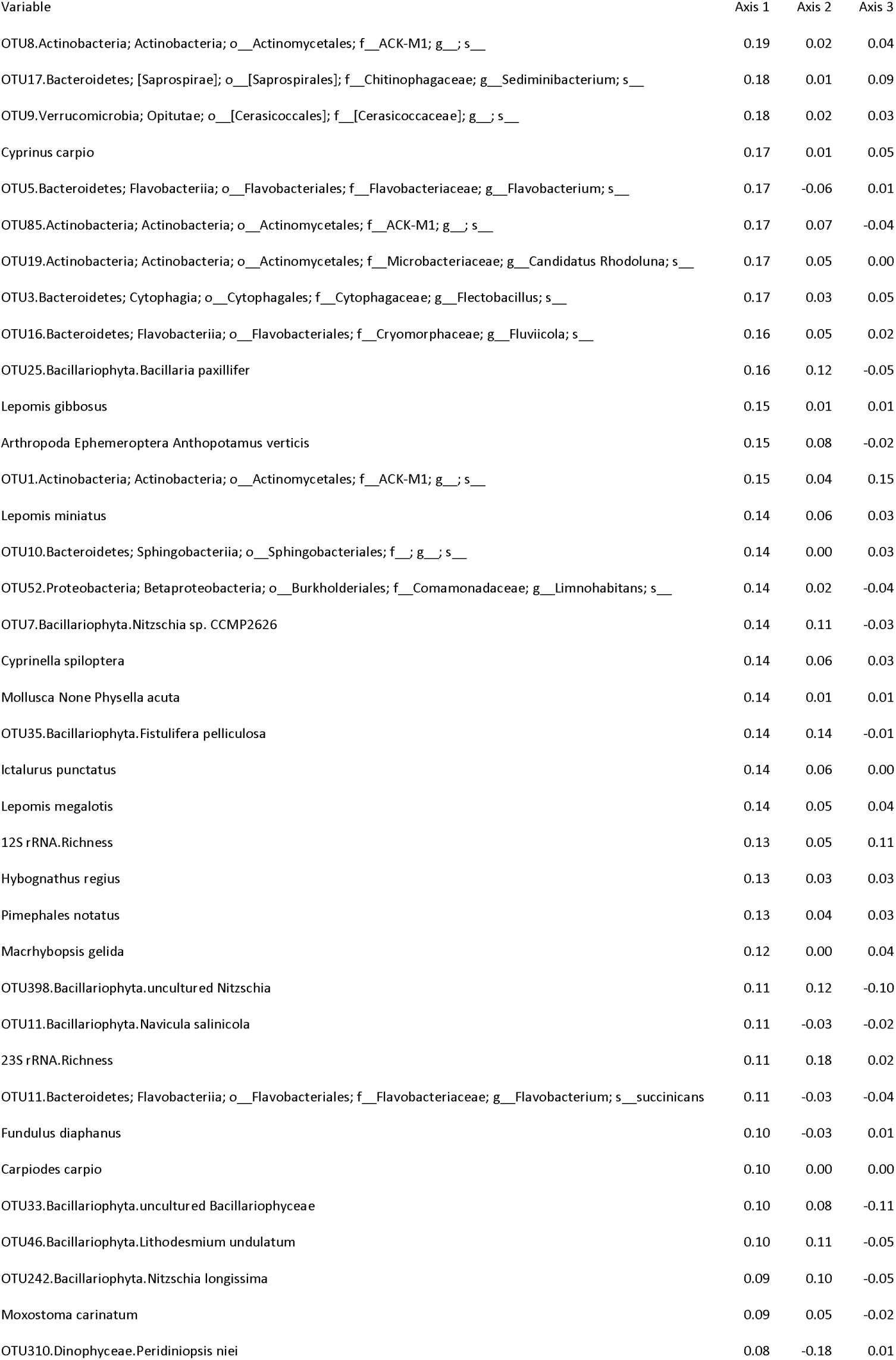
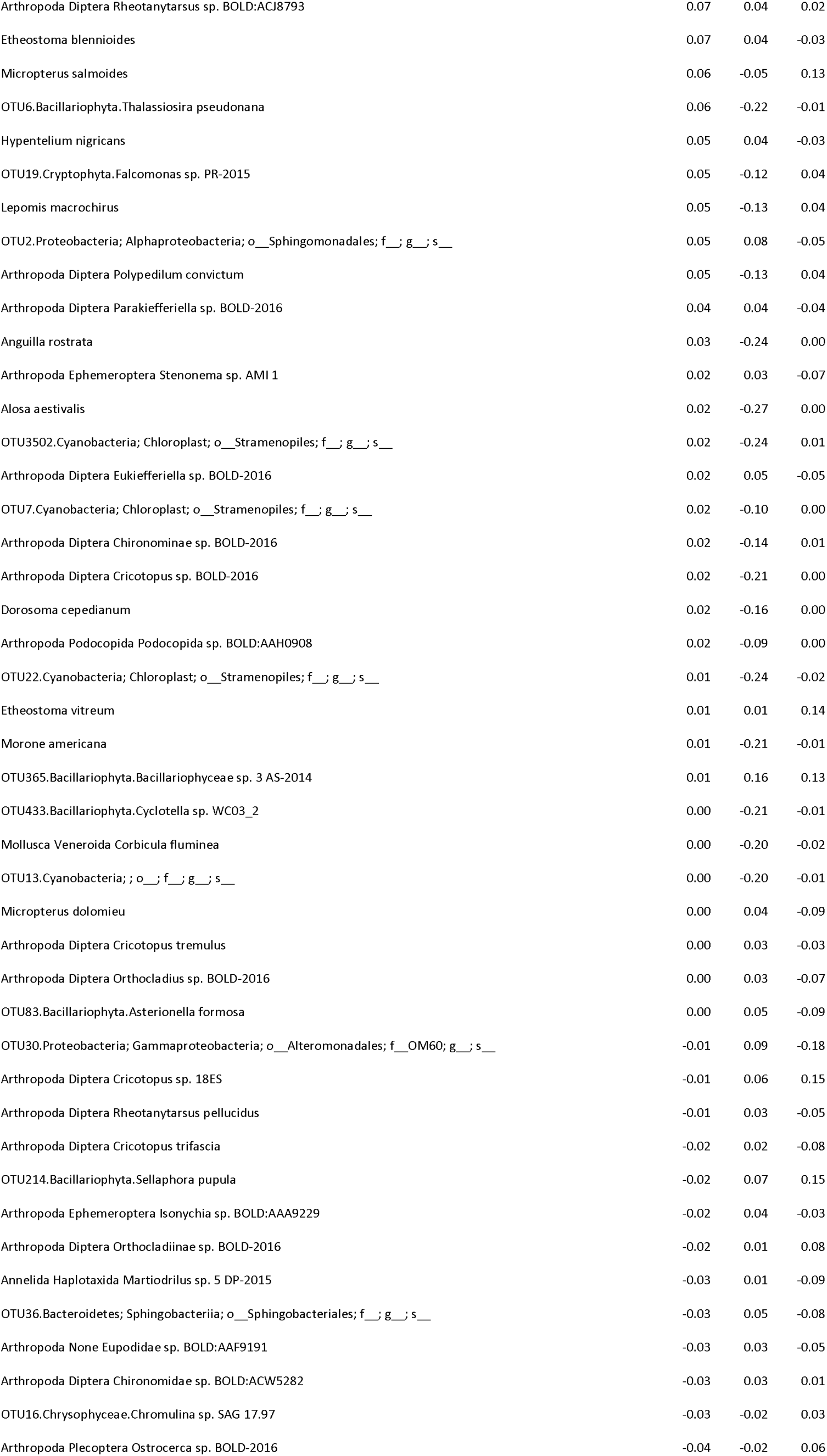
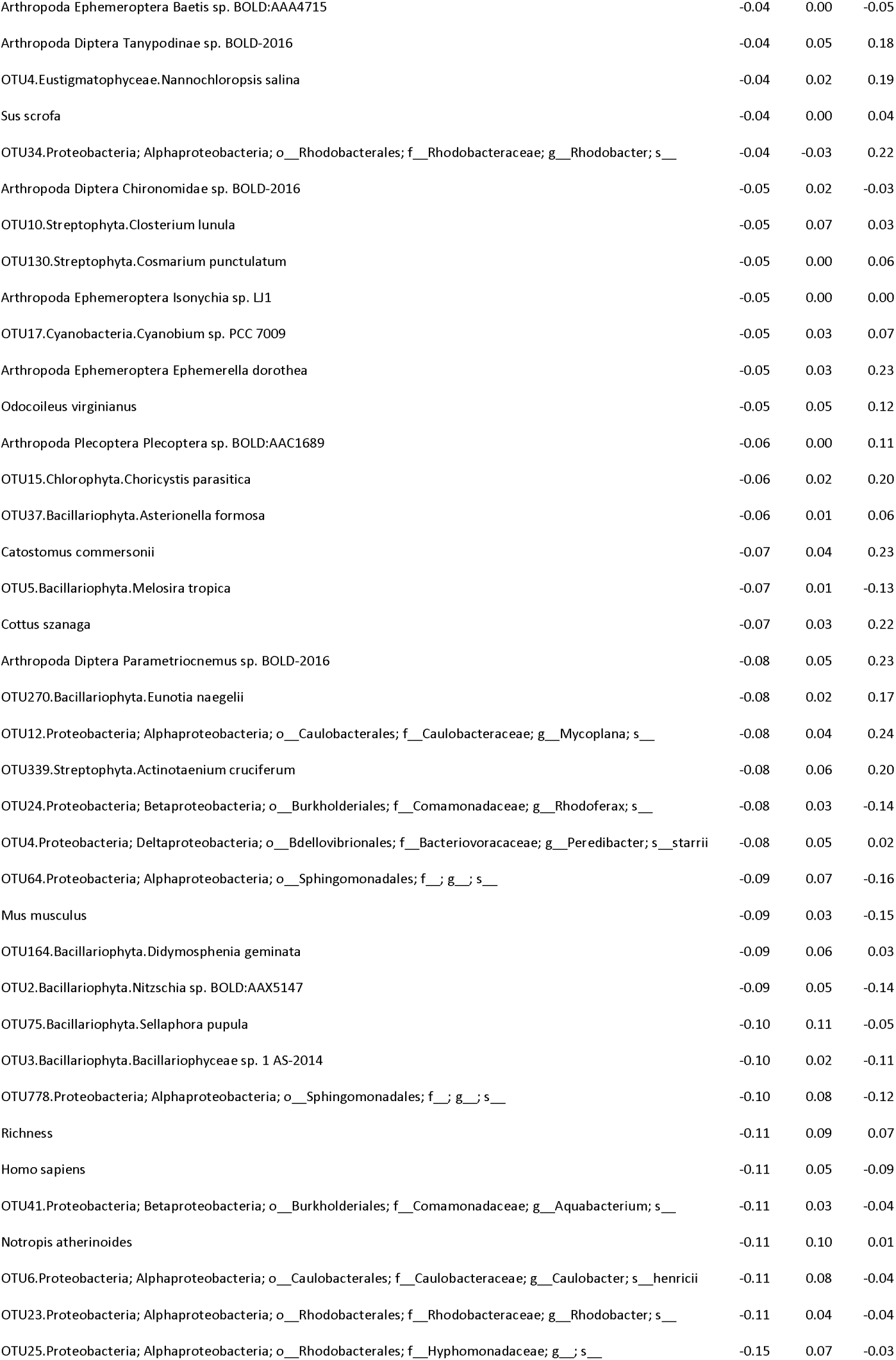

